# Quantifying rates of cell migration and cell proliferation in co-culture barrier assays reveals how skin and melanoma cells interact during melanoma spreading and invasion

**DOI:** 10.1101/124842

**Authors:** Parvathi Haridas, Catherine J. Penington, Jacqui A. McGovern, D. L. Sean McElwain, Matthew J. Simpson

## Abstract

Malignant spreading involves the migration of cancer cells amongst other native cell types. For example, *in vivo* melanoma invasion involves individual melanoma cells migrating through native skin, which is composed of several distinct subpopulations of cells. Here, we aim to quantify how interactions between melanoma and fibroblast cells affect the collective spreading of a heterogeneous population of these cells *in vitro*. We perform a suite of circular barrier assays that includes: (i) monoculture assays with fibroblast cells; (ii) monoculture assays with SK-MEL-28 melanoma cells; and (iii) a series of co-culture assays initiated with three different ratios of SK-MEL-28 melanoma cells and fibroblast cells. Using immunostaining, detailed cell density histograms are constructed to illustrate how the two subpopulations of cells are spatially arranged within the spreading heterogeneous population. Calibrating the solution of a continuum partial differential equation to the experimental results from the monoculture assays allows us to estimate the cell diffusivity and the cell proliferation rate for the melanoma and the fibroblast cells, separately. Using the parameter estimates from the monoculture assays, we then make a prediction of the spatial spreading in the co-culture assays. Results show that the parameter estimates obtained from the monoculture assays lead to a reasonably accurate prediction of the spatial arrangement of the two subpopulations in the co-culture assays. Overall, the spatial pattern of spreading of the melanoma cells and the fibroblast cells is very similar in monoculture and co-culture conditions. Therefore, we find no clear evidence of any interactions other than cell-to-cell contact and crowding effects.

## 1. Introduction

Melanoma is the deadliest form of skin cancer and arises due to the malignant transformation of melanocytes (Geller and Annas, 2003). In Australia, melanoma is reported to be the third most common cancer (Melanoma Institute Australia, 2016), and it is associated with high rates of mortality (Sneyd et al., 2013). However, if melanoma is detected early, before significant spreading occurs, prognosis after surgery is very good (Erdei et al., 2010; Faries and Ariyan, 2011). Therefore, understanding the mechanisms that drive melanoma spreading and invasion is very important.

Melanoma spreading takes place in a complex environment that including the extracellular matrix and many different kinds of cell types including: endothelial cells; keratinocytes; fibroblasts and immune cells (Cornil et al., 1991; Flach et al., 2011). Melanoma spreading in the dermis involves the movement of individual melanoma cells through an environment that also contains fibroblast cells (Li et al., 2007; Sriram et al., 2015). Previous experimental work suggests that melanoma cells can interact with fibroblast cells through diffusible factors, such as growth factors and cytokines, or by cell-to-cell contact and crowding (Flach et al., 2011; Goldstein et al., 2005; Labrousse et al., 2004; Ruiter et al., 2002; Zhou et al., 2015). In the experimental literature, these kinds of interactions are often broadly referred to as *cross-talk* between different subpopulations (Dvorankova et al., 2016; Ye et al., 2014). Although experimental studies indicate that fibroblasts can play a role in cancer progression, the precise details of how melanoma cells and fibroblast cells interact are not well understood (Kalluri and Zeisberg, 2006; Li et al., 2003).

Metastatic melanoma cells are known to grow in colonies, that are sometimes called nests (Baraldi et al., 2013; Schwartz et al., 2008). The spatial expansion of these colonies is driven by the rate at which individual melanoma cells move, and the rate at which individual melanoma cells proliferate. Therefore, to understand how quickly a population of melanoma cells spreads through the surrounding environment, it is important to develop techniques that allow us to quantify the rates of cell migration and cell proliferation (Treloar et al., 2013). Previous *in vitro* studies examining the spatial spreading of populations of melanoma cells have focused on monoculture experiments that contain only melanoma cells (Cornil et al., 1991; Im et al, 2012; Justus et al., 2014; Treloar et al., 2013). To make these kinds of *in vitro* studies more relevant to the *in vivo* environment, it is important to investigate, and quantify, how the spatial spreading of melanoma cells is affected by interactions with other cells types, such as fibroblasts.

In this study we perform a series of monoculture and co-culture barrier assays to examine the spatial and temporal patterns of the spreading of a heterogeneous cell population that is composed of both melanoma cells and primary fibroblast cells. All experiments in this work make use of the human metastatic melanoma cell line SK-MEL-28 (Fofaria and Srivastava, 2014), whereas the fibroblast cells are primary cells obtained from human donors. We first examine the spreading of melanoma cells and primary fibroblast cells separately, in a series of monoculture experiments. This allows us to quantify the rate of cell proliferation and the cell diffusivity for both melanoma cells and primary fibroblast cells, separately. Then, using our estimates of:

i. the melanoma cell diffusivity;
ii. the primary fibroblast cell diffusivity;
iii. the melanoma cell proliferation rate; and,
iv. the primary fibroblast cell proliferation rate,
 we investigate whether the solution of an appropriate mathematical model describing the co-culture experiments, parameterised using data from the monoculture experiments, is able to predict the patterns of spreading in a suite of co-culture experiments where both cell types are present in varying ratios. The procedure that we describe can be used to quantify the extent to which the interactions between the two cell types affect the co-culture experiments.

In summary, we present a method that can be used to identify potential interactions between two different cell types. In particular, we focus on interactions between primary fibroblast cells and SK-MEL-28 melanoma cells. Our hypothesis is that the rates at which these cells proliferate and migrate might be different when the cells are cultured in isolation to when the cells are cultured together. Overall, the results of our experimental and mathematical study supports the null hypothesis, since we find no clear evidence of any interactions other than cell-to-cell contact and crowding effects.

## 2. Experimental Methods

### 2.1 Melanoma Cell Culture

The metastatic melanoma cell line, SK-MEL-28, is cultured as described previously (Haridas et al., 2016). In brief, SK-MEL-28 melanoma cells are maintained in RPMI1640 medium (Thermo Scientific, Australia) supplemented with 10% fetal calf serum (FCS; Thermo Scientific), 2 mM L-glutamine (Thermo Scientific), 23 mM HEPES (Thermo Scientific), 50 U/ml of penicillin and 50 μg/ml of streptomycin (Thermo Scientific). The melanoma cell line is grown at 37 °C, in 5% CO_2_ and 95% air, and the cell line is routinely screened for *mycoplasma* contamination.

### 2.2 Primary fibroblast culture

Human skin discards are obtained from abdominoplasty and breast reduction surgeries (Xie et al., 2010). The epidermis is removed, discarded and the remaining dermis is used for fibroblast isolation. All experimental procedures are approved by the QUT research ethics committee (approval number QUT HREC #1300000063) and St Andrew’s Hospital ethics committee (approval number: Uniting Care Health 2003/46). The dermis is finely minced with a scalpel blade and placed in a 0.05% w/v collagenase A type I (Thermo Scientific) solution prepared in Dulbecco’s Modified Eagle’s Medium (DMEM) (Thermo Scientific) at 37 °C, in 5% CO_2_ and 95% air for 24 hours. The dermal cell solution is centrifuged at 212 g for 10 minutes, and cells are seeded into T75 cm^2^ flasks (Nunc^®^, Australia) in DMEM with 10% FCS, 2mM L-glutamine, 50 U/ml of penicillin and 50 μg/ml of streptomycin at 37 °C in 5% CO_2_ and 95% air.

### 2.3 Circular barrier assay

The spreading and proliferation of both primary fibroblast cells and SK-MEL-28 melanoma cells are examined using a two-dimensional circular barrier assay (Treloar et al., 2014a; Treloar et al. 2014b). Two types of experiments are performed. Firstly, in the monoculture experiments, the barrier assays are initialised with approximately 20,000 primary fibroblast cells (Fb monoculture), or approximately 20,000 SK-MEL-28 melanoma cells (SK monoculture). Secondly, in the co-culture experiments, assays are performed using three ratios of melanoma to fibroblast cells with the total number of initial cells held constant at approximately 20,000. We use three different ratios, and refer to these experiments as: co-culture 1; co-culture 2; and co-culture 3. Co-culture 1 experiments are initialised with approximately 15,000 primary fibroblast cells and approximately 5,000 SK-MEL-28 melanoma cells; co-culture 2 experiments are initialised with approximately 10,000 primary fibroblast cells and approximately 10,000 SK-MEL-28 melanoma cells; and, co-culture 3 experiments are initialised with approximately 5,000 primary fibroblast cells and approximately 15,000 SK-MEL-28 melanoma cells. We note that all experiments are initialised with approximately 20,000 cells in total. This means that the initial density of cells is less than half of the carrying capacity density, and this allows the cell populations to spread as a monolayer (Treloar et al. 2013; Treloar et al. 2014a) instead of piling up to form three-dimensional structures.

Clean and dried metal-silicone barriers, 6 mm in diameter (Aix Scientifics, Germany), are placed in a 24 well tissue culture plate (Nunc^®^) over glass coverslips (ProSciTech, Australia) containing 500 μl of full Green’s medium. The medium is made up of DMEM with Ham’s F12 medium (Thermo Scientific) in a 3:1 v/v ratio, 10% FCS, 2 mM L-glutamine, 50 U/ml of penicillin, 50 μg/ml of streptomycin, 180 mM adenine (Sigma Aldrich, Australia), 1 μg/ml insulin, 0.1 μg/ml cholera toxin (Sigma Aldrich), 0.01% non-essential amino acid solution (Thermo Scientific), 5 μg/ml transferrin (Sigma Aldrich), 0.2 μM triiodothyronine (Sigma Aldrich), 0.4 μg/ml hydrocortisone (Sigma Aldrich) and 10 ng/ml human recombinant epidermal growth factor (EGF; Thermo Scientific). The cell suspension is carefully pipetted into the barrier to ensure the cells are as evenly distributed as possible. Cells are allowed to attach to the plate for 2 hours in a humidified incubator at 37°C, 5% CO_2_ and 95% air, before the barriers are carefully removed (Treloar et al., 2013). The cell layer is washed with serum free medium (SFM; culture medium without FCS) and the cells are cultured in full Green’s medium. The culture plates are incubated at 37°C, 5% CO_2_ and 95% air for *t*=0, 24 and 48 hours. Each assay is performed in triplicates. Each assay is also repeated using primary fibroblast cells from three separate human donors.

### 2.4 Crystal violet staining

The cells grown on coverslips are washed with phosphate buffered saline (PBS; Thermo Scientific) and fixed for 20 minutes at room temperature using 10% neutral buffered formalin (United Biosciences, Australia), followed by staining the cells in 0.01% v/v crystal violet (Sigma Aldrich) in PBS. The coverslips are rinsed with PBS to remove excess stain and are air-dried. Images of the entire spreading cell population are acquired using a stereo microscope (Nikon SMZ 800) fitted with a Nikon digital camera.

### 2.5 Immunofluorescence

Cells grown on coverslips are fixed with 4% paraformaldehyde (Electron Microscopy Sciences, Australia) for 20 minutes at room temperature. Cell membranes are permeabilised with 0.1% v/v Triton X-100 (Merck Millipore, Australia) in PBS for 10 minutes, and the non-specific binding sites are blocked using 0.5% w/v bovine serum albumin (BSA) (Thermo Scientific) in PBS for 10 minutes. This is followed by the addition of primary antibody, *S100* in a ratio of 1:2000 (Dako, Australia) for an hour, and the secondary antibody *Alexa Fluor^®^* 555 in a ratio of 1:400 (Thermo Scientific) for an hour. Cells are washed with 0.5% BSA and the nuclei are stained with *dapi* in a ratio of 1:1000 (Sigma Aldrich) for 5 minutes. Coverslips are mounted on glass slides using ProLong^®^ Gold Antifade mountant (Thermo Scientific).

## 3. Mathematical modelling methods

### 3.1 Model summary

One way of providing further information about cancer progression is to interpret experimental observations using a mathematical model (Byrne, 2010). To quantify the role of various mechanisms acting in the monoculture and co-culture experiments we will use a continuum partial differential equation (PDE) model describing the collective spreading, proliferation and cell-to-cell crowding in a heterogeneous population of cells that is composed of two distinct subpopulations (Simpson et al. 2014). The PDE model is given by,

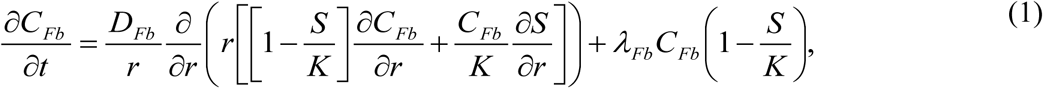

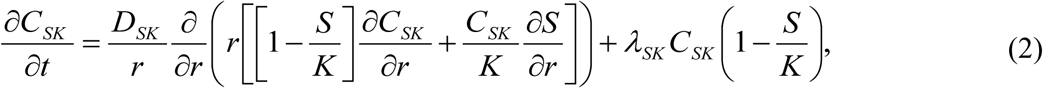
where *C_Fb_* (*r, t*) and *C_SK_* (*r, t*) are the density of fibroblast and melanoma cells, respectively, as a function of radial position, *r*, and time, *t*. The total cell density is given by *S*(*r, t*) = *C_Fb_* (*r, t*) + *C_SK_* (*r, t*), and the carrying capacity density is *K*. Since we consider circular barrier assays, in which the population of spreading cells always maintains a circular geometry over the entire duration of the experiment, Eq. (1)-(2) are written in terms of the radial coordinate, *r*, taking advantage of the axisymmetric geometry. Note that if there is just a single population present, Eq. (1)-(2) reduces to the standard Fisher-Kolmogorov equation in radial geometry (Treloar et al. 2014a).

### 3.2 Model parameters

There are five parameters in the co-culture model: (i) *D_Fb_* is the diffusivity of the primary fibroblast cells; (ii) *D_SK_* is the diffusivity of the SK-MEL-28 melanoma cells; (iii) *λ_Fb_* is the proliferation rate of the primary fibroblast cells; (iv) *λ_SK_* is the proliferation rate of the SK-MEL-28 melanoma cells; and (v) *K* is the carrying capacity density. Since the cells in our experiments spread as a monolayer, and the diameter of both the primary fibroblast cells and the SK-MEL-28 melanoma cells is approximately 20 μm (Treloar et al. 2013; Treloar et al. 2014a), we estimate the carrying capacity density by assuming that the maximum density of cells corresponds to hexagonal packing of disks of diameter 20 μm, giving *K* = 2.8×10^−3^ cells/μm^2^. With this assumption there are now four unknown parameters in Eq. (1)-(2).

### 3.3 Initial condition

The PDE model can be used to simulate both co-culture and a monoculture barrier assays. To simulate a co-culture assay we specify appropriate non-zero initial conditions for both *C_Fb_* (*r*,0) and *C_SK_* (*r*,0), chosen to match the initial cell density in the co-culture experiments. Alternatively, to simulate a fibroblast monoculture assay, we set *C_SK_* (*r*,0) = 0 and specify some appropriate non-zero initial condition for *C_Fb_*(*r*,0). Similarly, to simulate a melanoma monoculture assay we set *C_Fb_* (*r*,0) = 0 and specify some appropriate non-zero initial condition for *C_SK_* (*r*,0).

### 3.4 Numerical solution

Regardless of whether we use the PDE model to simulate a monoculture or co-culture assay, we always solve Eq.(1)-(2) numerically. Spatial derivatives are approximated using a central difference approximation on a uniformly-spaced finite difference mesh, with mesh spacing Δ*r* The resulting system of coupled nonlinear ordinary differential equations is solved using a backward Euler approximation, with time steps of duration Δ*t*. The nonlinear ordinary differential equations are linearised using Picard iteration with a convergence tolerance of *ε* (Chapra and Canale, 1998).

### 3.5 Model background

Before applying Eq. (1)-(2) to our experimental data set, it is useful to briefly explain the origin of the PDE model and the underlying assumptions. The model was described and presented by us previously (Simpson et al. 2014). In that previous work we consider both a stochastic random walk process and the associated continuum limit PDE description. In brief, the lattice based random walk model describes the collective motion of a population of two potentially distinct subpopulations of cells. Cells in both subpopulations undergo nearest neighbour motility events, where cells attempt to step a distance of Δ, at some specified constant rate. Here, Δ corresponds to the average cell diameter. Potential motility events are unbiased so that the direction of movement is chosen with equal probability. Crowding effects are incorporated by ensuring that any potential motility event that would place a cell on an occupied site is aborted. The discrete model also allows cells to proliferate, at some other specified constant rate. A potential proliferation event will involve a cell placing a daughter cell, of the same subpopulation, on a randomly chosen nearest neighbour lattice site. Again, crowding effects are incorporated by ensuring that any potential proliferation event that would place a daughter cell on an occupied lattice site is aborted. The continuum limit description of this discrete model, in a radially symmetric geometry, is Eq. (1)-(2) (Simpson et al., 2014).

The system of PDEs, given by Eq. (1)-(2), corresponds to a coarse-grained description of the cell-to-cell crowding effects that are explicitly described in the discrete random walk model. For example, the nonlinear diffusion terms in Eq. (1)-(2) correspond to hard-core exclusion in the motility mechanism of the discrete model. Similarly, the nonlinear source terms in Eq. (1)-(2) correspond to the proliferation mechanism of the discrete model.

## 4. Experimental results and discussion

### 4.1 Diameter of the spreading population

We first investigate the spatial expansion of the cell populations over time. Results in Fig. 1(a)-(i) show the spreading populations from *t*=0 until *t*=48 hours. To quantify the spatial spreading, we calculate the diameter of each spreading population at *t*=0, 24 and 48 hours. To achieve this we use ImageJ (2016) to automatically detect the position of the leading edge of the spreading population using the Sobel method (Treloar and Simpson, 2013; Johnston et al. 2014). ImageJ also provides an estimate of the area contained within the leading edge of the spreading population. Using this estimate of area, we assume that the spreading population remains approximately circular, allowing us to convert the area estimate into an estimate of the equivalent circular diameter. The diameter of the spreading populations is shown in Fig. 1(j) where we see that there is an increase in the diameter with time in all cases. However, we observe that the rate of increase in the diameter in some experiments is different. For example, we observe that Fb monoculture experiments spread fastest, whereas the SK monoculture experiments spread slowest. In comparison, the co-culture experiments spread at an intermediate rate.

**Fig. 1.**
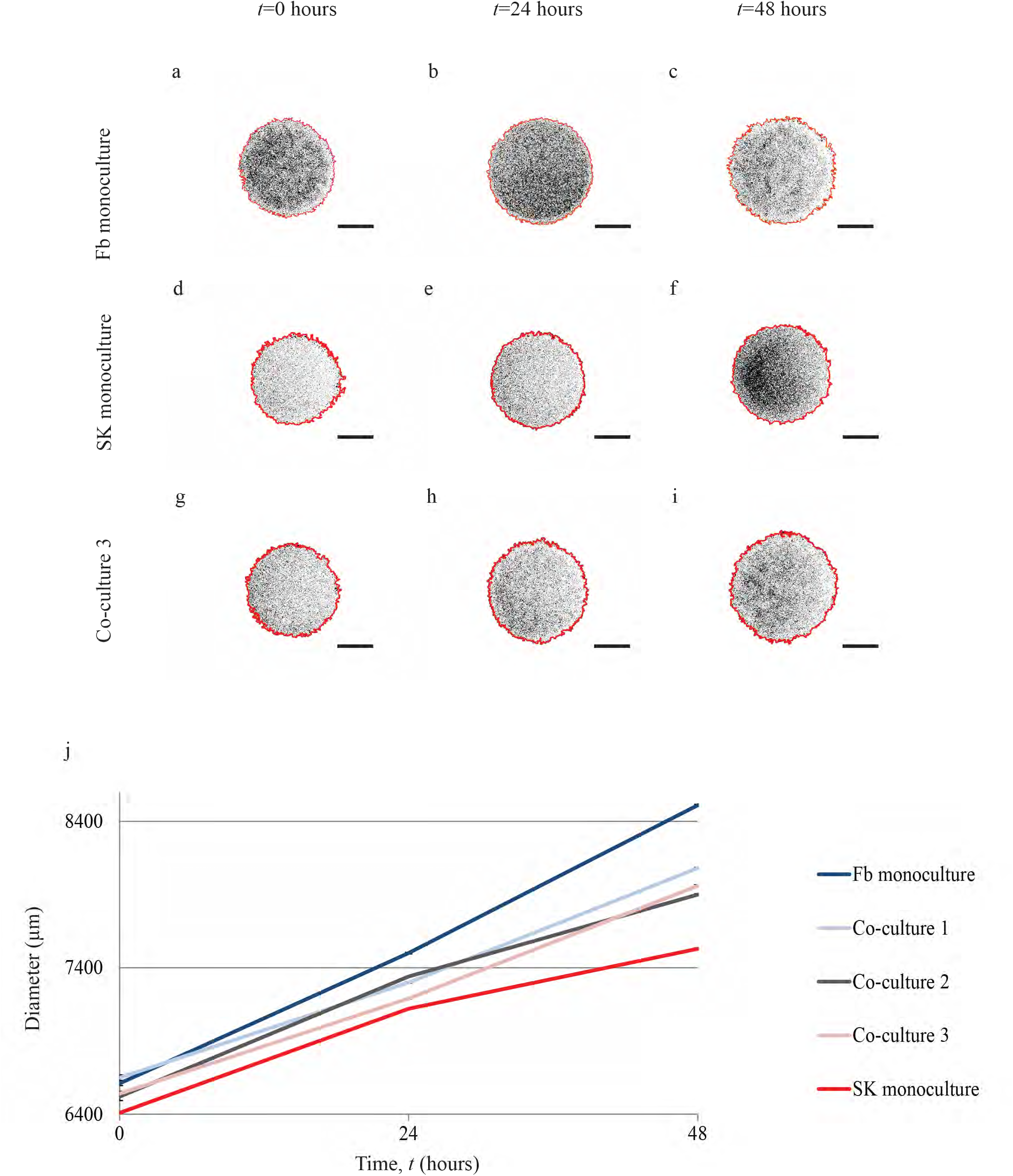
Spatial spreading of cell populations over 48 hours. Images in (a)-(c) show monoculture barrier assays initialised with approximately 20,000 primary fibroblasts (Fb monoculture), (d)-(f) show monoculture barrier assays initialised with approximately 20,000 SK-MEL-28 melanoma cells (SK monoculture), and (g)-(i) show co-culture barrier assays initialised with approximately 10,000 SK-MEL-28 melanoma cells and approximately 10,000 primary fibroblast cells (co-culture 2). Images show the spreading of the population at *t*=*0*, 24 and 48 hours, as indicated. The red outline shows the position of the leading edge detected using ImageJ. Each scale bar is 3000 μm. Data in (j) show the increase in average diameter of the spreading cell populations with time (n=3). Each initial ratio of cells is shown using a different colour, as indicated. Co-culture 1 corresponds to experiments initialised with approximately 15,000 primary fibroblasts and approximately 5,000 SK-MEL-28 melanoma cells, co-culture 2 corresponds to experiments initialised with approximately 10,000 primary fibroblasts and approximately 10,000 SK-MEL-28 melanoma cells, and co-culture 3 corresponds to experiments initialised with approximately 5,000 primary fibroblasts and approximately 15,000 SK-MEL-28 melanoma cells.

Although these results focusing on the rate at which the leading edge of the populations spread is insightful, they do not provide any information about how the primary fibroblast cells and the SK-MEL-28 melanoma cells are distributed throughout the spreading populations. To provide this additional information, we use a more sophisticated experimental approach.

### 4.2 Cell type identification in co-cultures

To extend our initial investigation about the spatial expansion of the cell populations, we quantify the spatial distribution of primary fibroblast cells and SK-MEL-28 melanoma cells throughout the spreading populations. To achieve this we must distinguish the primary fibroblast cells from the SK-MEL-28 melanoma cells within these heterogeneous populations. The metastatic melanoma cell line, SK-MEL-28 can be reliably and exclusively identified using the *S100* marker (Haridas et al., 2016). However, it is challenging to identify primary fibroblast cells in a heterogeneous population because many because fibroblast markers, like *vimentin* and *alpha smooth muscle actin*, are also expressed by other migrating cell types present in the population (Kalluri and Zeisberg, 2006; Marsh et al., 2013; Sugimoto et al., 2006). To deal with this complication we use *dapi* to stain the cell nuclei of all cells, capturing both the SK-MEL-28 melanoma cells and the primary fibroblast cells. By counting the number of *dapi*-positive cells and subtracting the number of *S100* positive cells, we are able to reliably estimate the number of primary fibroblast cells in each image.

Images showing the entire spreading populations are superimposed with an immunostained transect that passes through the centre of the cell population in Fig. 2. These immunostained transects allow us to extract detailed information about the spatial distribution of primary fibroblast cells and SK-MEL-28 melanoma cells in each experiment. Greyscale images showing the entire spreading population at *t*=24 and 48 hours are shown in Fig. 2(a), (c), (e), (g), (i), (k), (m), (o), (q) and (s). The central region of the transect, as indicated, is magnified and shown in Fig. 2(b), (d), (f), (h), (j), (l), (n), (p), (r) and (t). Our results in Fig. 2 show that we are able to clearly and reliably identify the two different cell types in the experiments. We are confident in our results because there are no *S100* positive cells in the primary fibroblast monoculture experiment (Fig. 2(b), (d)), and we observe an increasing proportion of *S100* positive cells in co-culture 3 (Fig. 2(m)-(p)), compared to co-culture 2 (Fig. 2(i)-(l)). Similarly, we observe an increasing proportion of *S100* positive cells in co-culture 2 (Fig. 2(i)-(l)), compared to co-culture 1 (Fig. 2(e)-(h)).

**Fig. 2.**
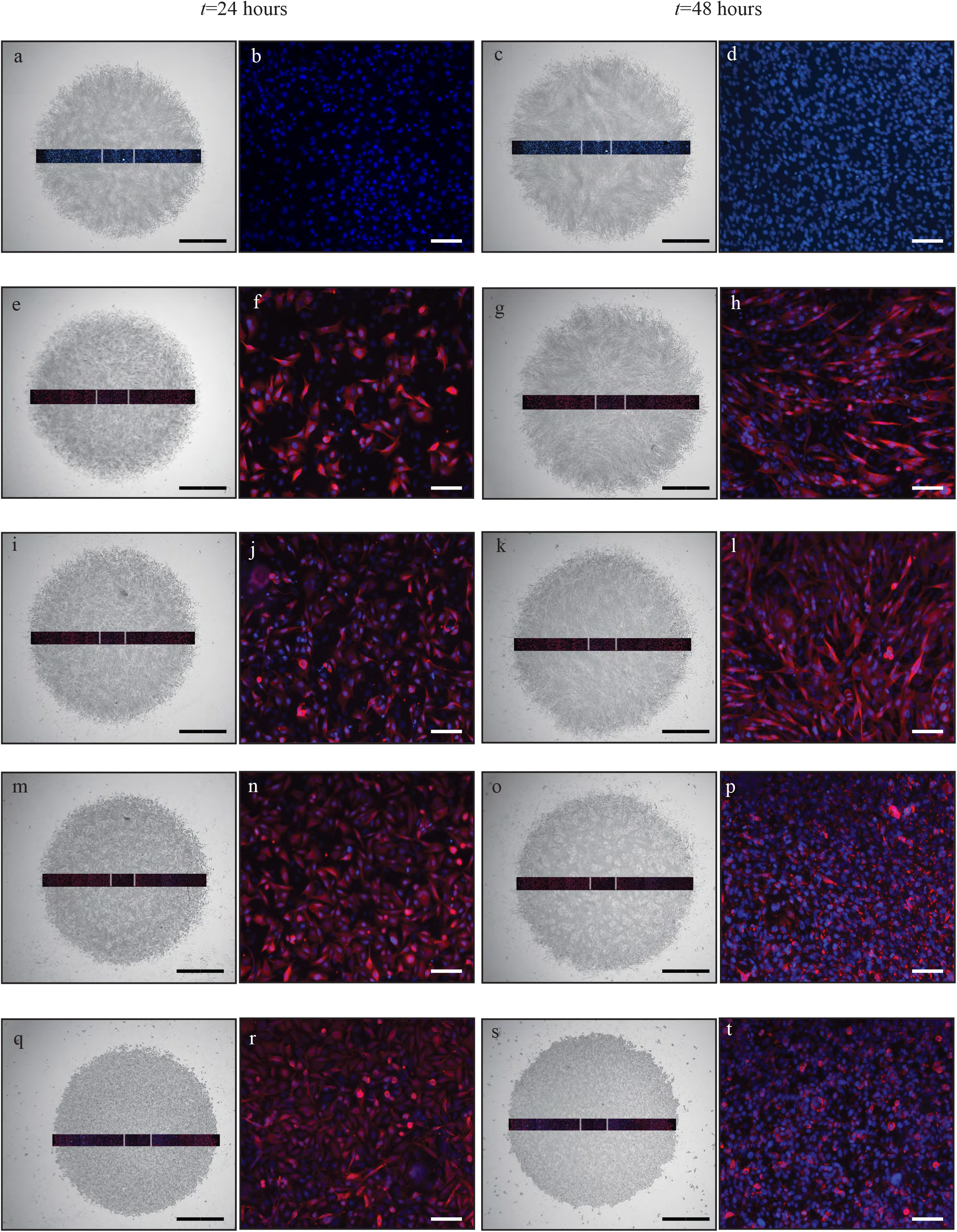
Experimental images of spreading cell populations and corresponding immunofluorescence images to detect the composition of the co-culture assays, at *t*=*24* and 48 hours. The two left-most columns of images correspond to *t*=24 hours, and the two right-most columns of images correspond to *t*=48 hours. Results in (a) and (c) correspond to Fb monoculture; (e) and (g) correspond to co-culture 1; (i) and (k) correspond to co-culture 2; (m) and (o) correspond to co-culture 3; and (q) and (s) correspond to SK monoculture, as indicated. A transect showing immunofluorescence staining is superimposed on each greyscale image, and the transect passes through the centre of each spreading population. The vertical white lines on each transect indicates the central region of the transect, and the central regions in (a), (c), (e), (g), (i), (k), (m), (o), (q) and (s) are magnified, and shown in (b), (d), (f), (h), (j), (l), (n), (p), (r) and (t), respectively. In the immunofluorescence images, all cell nuclei (Fb + SK) are stained with *dapi* (blue), whereas just the SK-MEL-28 melanoma cells (SK) are stained with *S100* (red). The scale bar in all greyscale images is 2000 μm, and the scale bar in all immunofluorescence images is 100 μm.

### 4.3 Construction of cell density histograms

To quantify how the two subpopulations of cells are spatially distributed within the heterogeneous spreading populations, we construct cell density histograms. To do this, we count the number of cells in many equally spaced subregions across each transect, as shown in Fig. 3(a). We use immunofluorescence to identify primary fibroblast cells and SK-MEL-28 melanoma cells as described in Section *4.2*, and as shown in Fig. 3(b)-(d). An estimate of the cell density for each cell type along the transect is calculated by counting the number of primary fibroblast cells and the number of SK-MEL-28 melanoma cells in each subregion, and dividing these numbers by the area of the subregion. A histogram showing cell density as a function of position is generated for each experimental replicate in each experimental condition. Averaging the histograms from each experimental replicate gives an averaged histogram, as shown in Fig. 3(e). The raw data showing the cell density information for each experimental replicate is also provided (Supplementary Material-1).

**Fig. 3.**
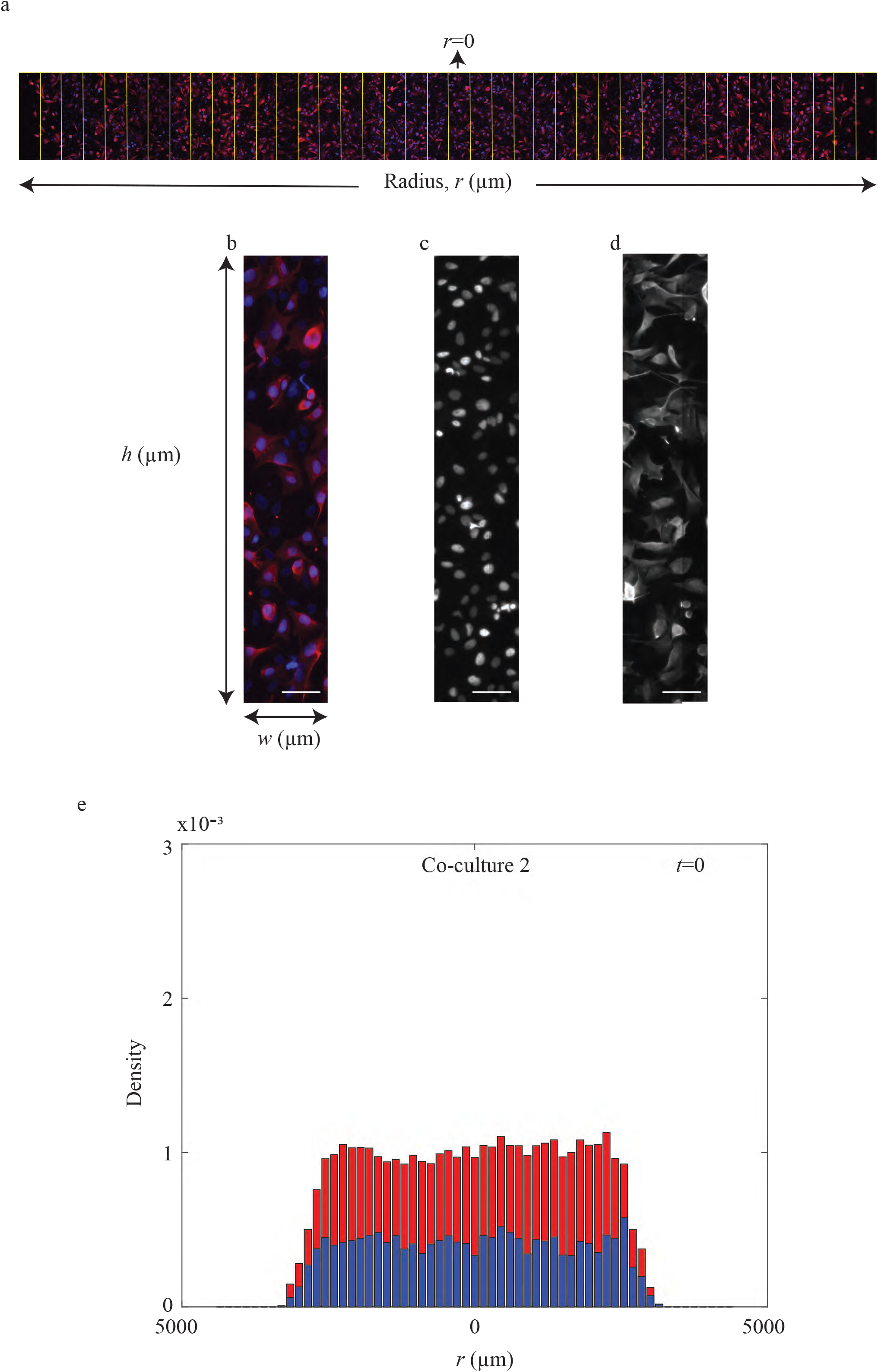
Immunofluorescence staining identifies the spatial and temporal patterns of cell spreading in the co-culture barrier assays. (a) Example transect through the centre of a spreading population. The centre of the population corresponds to *r*=0, and the distance from the centre of the population is measured by the radial coordinate, *r*>0. Each transect is divided into many equally-spaced subregions, each of width *w*=150 μm and height *h*. The value of *w* is constant, fixed at 150 μm in all experiments. However, the height of the subregion, *h*, varied between 622-817 μm in different experiments, but the height is constant each transect in each particular experiment. To quantify the density of cells across the transect, we count the number of cells of each type in each subregion, and divide by the area of the subregion to give an estimate of the density of each cell type, at each radial position, *r*. To count the number of cells in each subregion we use immunofluorescence staining, as shown in (b). The staining in (c) shows *dapi* staining (Fb + SK), whereas the staining in (d) shows *S100* (SK) staining. The number of primary fibroblast cells in each subregion is the difference between the total number of *dapi*-positive nuclei and the number of *S100*-positive cells in each subregion. The scale bar in (b)-(d) is 100 μm. Using these cell counts, we construct the cell density histogram, as shown in (e), illustrating the spatial variation in cell density at *t*=0 in an experiment corresponding to co-culture 2. The blue section in the histogram shows the density of primary fibroblast cells, the red section shows the density of SK-MEL-28 melanoma cells, and the total height of the histogram shows the total cell density.

A series of averaged cell density histograms at *t*=0, 24 and 48 hours are shown in Fig. 4. Each histogram shows the average density of cells across the entire transect. The radial position is given by *r*>0. The centre of the spreading cell population corresponds to *r*=0, and the population spreads in both directions, away from the centre. The blue section in the histograms indicate the average density of primary fibroblast cells, the red section shows the average density of SK-MEL-28 melanoma cells, and the total height of the histogram shows the average total cell density.

**Fig. 4.**
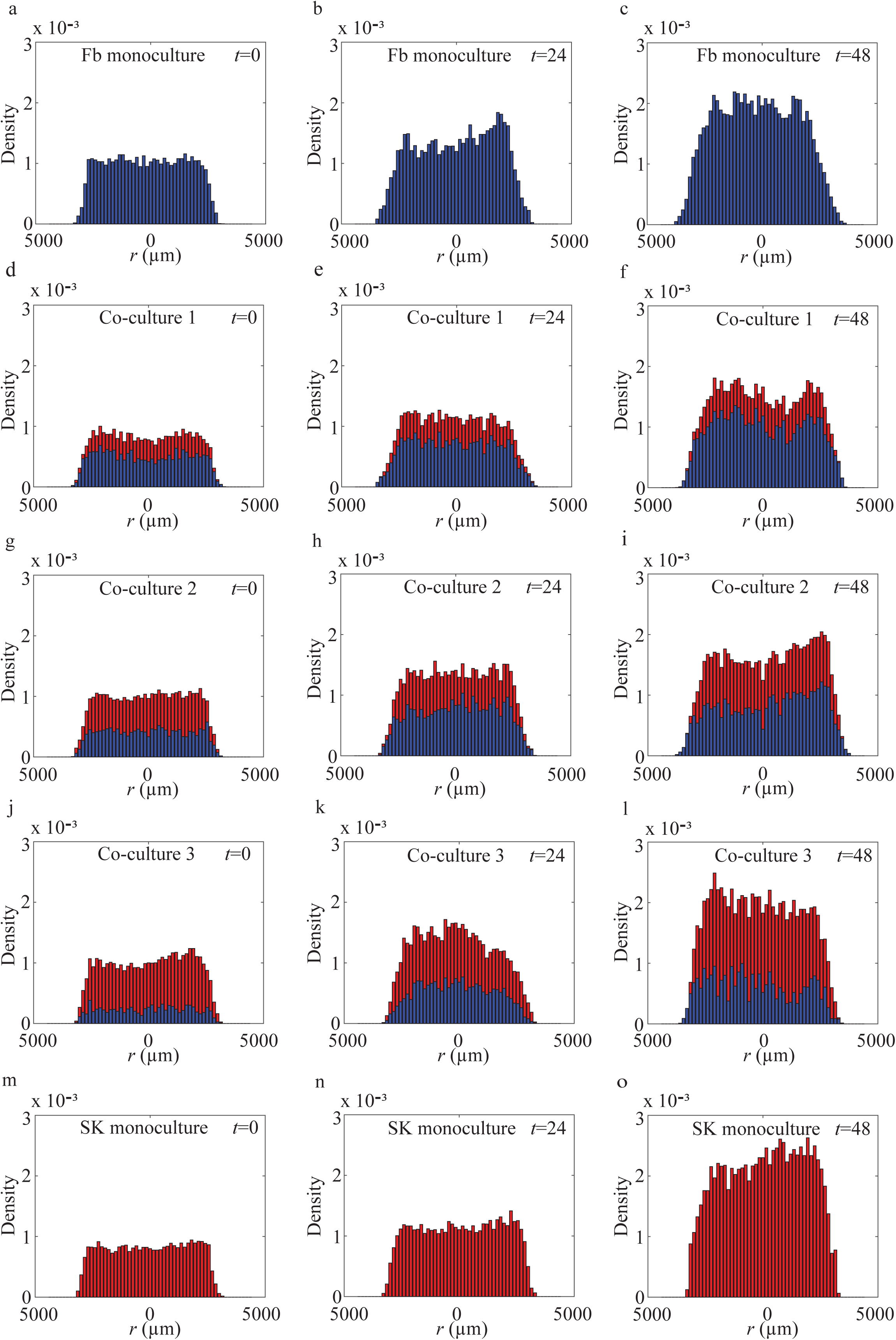
Summary of cell density profiles. Cell density histograms for individual experiments, constructed using the technique presented in Fig. 3, are averaged across three identically prepared experimental replicates (n=3) to give a series of averaged cell density histograms. Averaged cell density histograms in (a)-(c) correspond to Fb monoculture; (d)-(f) correspond to co-culture 1; (g)-(i) correspond to co-culture 2; (j)-(l) correspond to co-culture 3; and (m)-(o) correspond to SK monoculture, at *t*=0, 24 and 48 hours, as indicated. The blue section in the histogram shows the density of primary fibroblast cells, the red section shows the density of SK-MEL-28 melanoma cells, and the total height of the histogram shows the total cell density.

Results in Fig. 4(a), (d), (g), (j) and (m) show the histograms at *t*=0. These results confirm that all barrier assays are initialised such that the total population of cells is approximately uniformly distributed across the transect, with a total cell density of approximately 1×10^−3^ cells/μm^2^, which is less than half the carrying capacity density of these cells in a monolayer (Supplementary Material-2). However, the ratio of melanoma to primary fibroblast cells differs in Fig. 4(a), (d), (g), (j) and (m). For example, the profile in Fig. 4(a) contains only primary fibroblast cells, the prolife in Fig. 4(m) contains only melanoma cells, and the profiles in Fig. 4(d), (g) and (j) contain both melanoma and primary fibroblast cells with an increasing ratio of melanoma to fibroblast cells, respectively.

Our results in each row in Fig. 4 show how the cell density changes with time. The middle column of results corresponds to *t*=24 hours, and the right-most column corresponds to *t*=48 hours. Comparing results in Fig. 4(c) and Fig. 4(o) shows that the cells in the Fb monoculture experiments spread further than the cells in the SK monoculture experiments, and result this is consistent with the leading edge results in Fig. 1. Furthermore, comparing the shape of the cell density histograms in Fig. 4(c) and Fig. 4(o) shows that the leading edge of the cell density profile is sharper in the SK monoculture experiments than for the Fb monoculture experiments. While we observe differences in the rate of the spatial extent of the spreading of the two monoculture experiments, we observe that the increase in cell density towards the centre of the population, at *r*=0, is very similar. For example, at *t*=*0* the cell density at the centre of both monoculture experiments is approximately 1×10^−3^ cells/μm^2^, and after 48 hours the cell density at the centre of both monoculture experiments has approximately doubled to a density of 2×10^−3^-2.5×10^−3^cells/μm^2^.

Comparing the time evolution of the cell density patterns in the co-culture experiments with the monoculture experiments suggests that there are minimal differences in the behaviour of the monoculture and co-culture experiments. For example, co-culture 1 that is initiated with approximately 15,000 primary fibroblast cells and approximately 5,000 SK-MEL-28 melanoma cells behaves in a very similar way to the Fb monoculture experiment in terms of both the spatial extent of the spreading of the total population and the total increase in the cell density towards the centre of the spreading population. Similarly, co-culture 3, that is initiated with approximately 5,000 primary fibroblast cells and approximately 15,000 SK-MEL-28 melanoma cells behaves in a very similar way unto the SK monoculture experiment in terms of both the spatial extent of the spreading of the total population and the total increase in the cell density towards the centre of the spreading population. Results for co-culture 2 lie between co-culture 1 and co-culture 3. An interesting feature of co-culture 1 and co-culture 2 is that at both *t*=24 and *t*=48 hours, we see that the primary fibroblast cells dominate the total population right at the leading edge of the heterogeneous population of cells. This is consistent with our observation that the primary fibroblast cells spread faster than the SK-MEL-28 melanoma cells in the monoculture experiments.

Now that we have presented, and discussed, the cell density histograms for the monoculture and co-culture experiments, we will further explore the similarities and differences between the experiments by calibrating a mathematical model to these data. Combining our experimental results with a mathematical model will allow us to explore, in more detail, the question of whether the primary fibroblast cells and/or the SK-MEL-28 melanoma cells behave differently when grown in monoculture or in co-culture conditions.

## 5. Mathematical results and discussion

### 5.1 Estimating parameters for the monoculture experiments

Since fibroblast cells and melanoma cells are cultured separately in the monoculture experiments, the coupled model, given by Eq. (1)-(2), uncouples to give

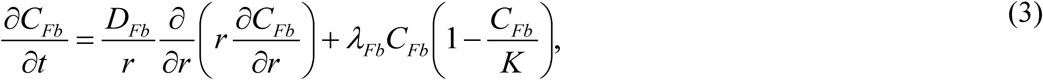

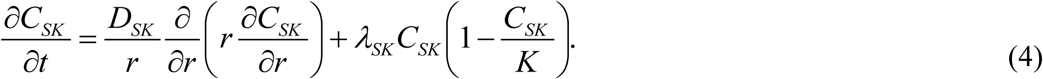

Equations (3)-(4) are two, uncoupled, single-species Fisher-Kolmogorov equations in radial coordinates that we will use to describe the Fb monoculture and SK monoculture experiments, respectively. There are four parameters to be estimated: *D_Fb_*; *D_SK_*; *λ_Fb_* and *λ*_SK_. We now explain how these parameters can be estimated separately, using data in Fig. 4(a)-(c) for the Fb monoculture experiment, and using data in Fig. 4(m)-(o) for the SK monoculture experiment. Following the approach of Johnston et al. (2015), we note that in the central region of the monoculture experiments, where *r* <1425 μm, the cell density profile is approximately spatially uniform in both the fibroblast monoculture and the melanoma monoculture (Fig. 4). This region approximately corresponds to the middle third of the spreading population, and hence this region is well away from the leading edge of the spreading populations. Since, locally in the centre of the fibroblast monoculture experiment we have *∂C_Fb_* / *∂r* ≈ 0, and locally in the centre of the melanoma monoculture experiment we have *∂C_SK_* / *∂r* ≈ 0, the two uncoupled PDEs, Eq. (3)-(4), simplify to two uncoupled ordinary differential equations (ODE) that can be written as

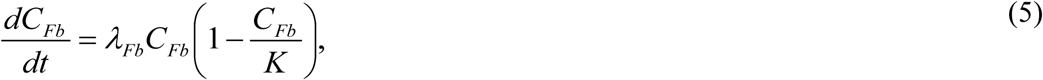

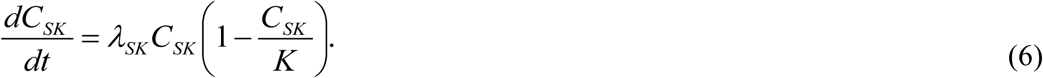
The solutions of Eq. (5)-(6) are

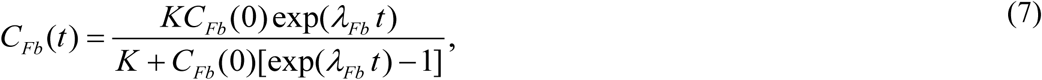

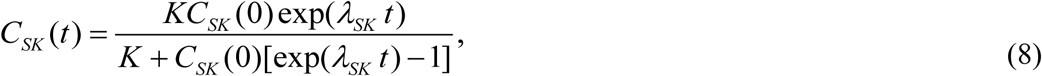
where *C_Fb_* (0) is the initial density of the fibroblast cells in the central region of the Fb monoculture experiments, and *C_SK_* (0) is the initial density of the melanoma cells in the central region of the SK monoculture experiments. Estimates of *λ_Fb_* and *λ_SK_* are obtained by choosing these parameters so that *C_Fb_* (*t*) and *C_SK_* (*t*), given by Eq. (7)-(8), match the experimental data from the central region of the fibroblast monoculture experiments and the melanoma monoculture experiments, respectively (Supplementary Material-2). In summary, matching these solutions to our experimental data gives us a range of estimates: 0.02 ≤ 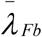 ≤ 0.04/hour and 0.03 ≤ 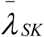 ≤ 0.05 /hour. Here we use the overbar notation to indicate the least-squares estimates of the parameters, and the range of estimates corresponds to the sample mean plus or minus one sample standard deviation calculated using the three identically prepared experimental replicates of the monoculture experiments. It is interesting to note that these estimates of the proliferation rate for the melanoma cells and the fibroblast cells are approximately equal. Furthermore, the proliferation rates correspond to a doubling time of approximately 23 hours, and this is consistent with previous results (Treloar et al. 2013).

Given our estimates of *λ_Fb_* and *λ_SK_*, we solve Eq. (3)-(4) across the entire domain, 0 < r < 4350 μm, and match the numerical solution of each uncoupled PDE with the averaged cell density profiles across the entire domain for both monoculture experiments. Setting the proliferation rates to be in the middle of the range previously identified (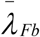 = 0.03/hour and 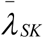 = 0.04/hour), we obtain estimates of 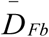 ≈ 1200 μm^2^/hour, and 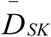 ≈ 170 μm^2^/hour (Supplementary Material-2). Unlike our estimates of the proliferation rates, the estimate of the cell diffusivity for the fibroblast cells is an order of magnitude higher than the estimate of the cell diffusivity for the melanoma cells. Our estimate for the cell diffusivity of the human primary fibroblast cells is very similar to previous estimates of the cell diffusivity for 3T3 mouse fibroblast cells, which have been reported to be approximately 800-2900 μm^2^/hour (Treloar et al. 2014a). Furthermore, our estimate of the cell diffusivity for the SK-MEL-28 melanoma cells is similar to previous estimates for other metastatic melanoma cell lines, which have been reported to be approximately 160-250 μm^2^/hour (Treloar et al. 2013).

### 5.2 Predicting collective cell spreading in the co-culture experiments

Given our parameter estimates from the monoculture experiments, we are interested to investigate whether the solution of the coupled co-culture model, Eq. (1)-(2), can accurately predict the spatial and temporal patterns of spreading in the co-culture experiments when parameterised in the same way. Examining this question will provide insight into whether the fibroblast and/or melanoma cells behave differently in monoculture than they do in co-culture. In summary, if we can find a unique choice of *D_Fb_*; *D_SK_*; *λ_Fb_* and *λ_SK_* for which:

i. the solution of the coupled system, Eq. (1)-(2), matches the experimental data for all three co-culture assays;
ii. the solution of Eq. (3) matches the experimental data for the fibroblast monoculture assay; and
iii. the solution of Eq. (4) matches the experimental data for the melanoma monoculture assay,
 it would be reasonable to conclude that the migratory and proliferative behaviour of the melanoma and fibroblast cells appears to be independent of whether these two cell types are cultured separately or together. In contrast, if we must choose very different parameter values to match the monoculture experiments compared to the parameter values required to match the co-culture experiments, then our results would suggest that the cells behave very differently in monoculture and co-culture environments.

Results in Fig. 5 compare the spatial and temporal evolution of the two monoculture assays and the three co-culture assays together with the solution of the appropriately parameterised mathematical models. Using our parameter estimates from the monoculture experiments as an initial estimate, we manually adjusted the parameters and find that setting *D_Fb_* ≈ 1200 μm^2^/hour; *D_SK_* ≈ 170 μm^2^/hour; *λ_Fb_* = 0.03/hour and *λ_SK_* = 0.03 /hour leads to a reasonably accurate match across both the two monoculture assays and the three co-culture assays (Supplementary Material-2). Given that we are able to match both the monoculture and co-culture experiments using a single combination of parameters, this suggests that the only interactions necessary to explain the experimental observations are cell-to-cell contact and crowding effects. In particular, no additional cross-talk mechanisms, such as interactions mediated by the production of signalling factors, is required to explain our experimental observations.

**Fig. 5.**
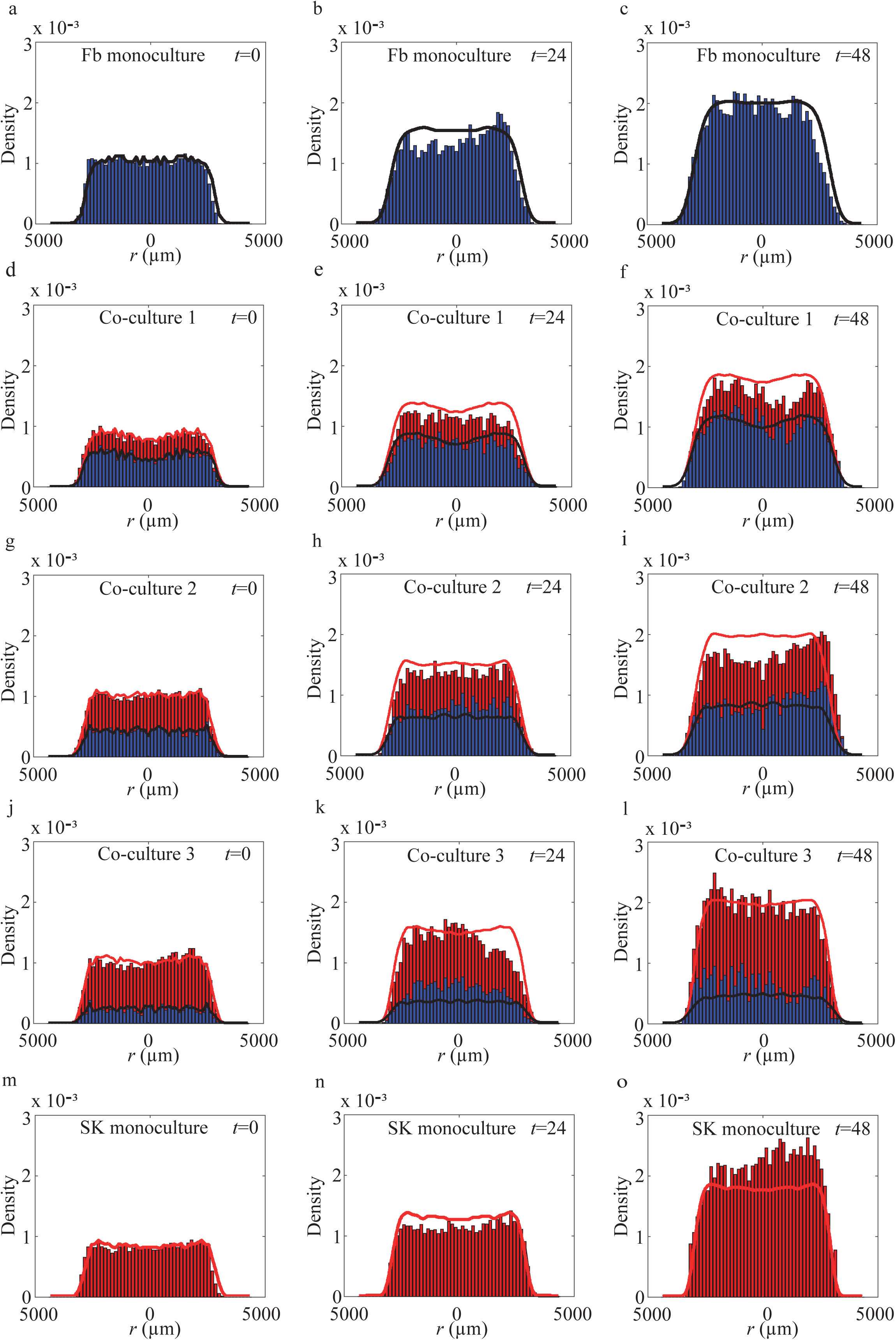
Comparison of average cell density profiles and the solution of the mathematical model, Eq. (1)-(2). The experimental data is presented in the same format as presented in Fig. 4. All experimental data are superimposed with appropriate numerical solutions of Eq. (1)-(2), with *C_Fb_* (*r, t*) shown in black, and *S* (*r, t*) shown in red, and *C_SK_* (*r, t*) is the difference between the red curve and the black curve. The initial condition for *C_Fb_* (*r*,0) and *S*(*r*,0), shown in (a), (d), (g), (j) and (m), are chosen to match the observed experimental data at *t*=0. The parameters used to solve Eq. (1)-(2) are: *D_Fb_* = 1200 μm^2^/hour; *D_SK_* = 170 μm^2^/hour; *λ_Fb_* = 0.03 hour^−1^; *λ_SK_* = 0.03 hour^−1^ and *K* = 2.8 × 10^−3^ cells/μm^2^. The equations are solved on 0 < *r* < 4350μm. Zero flux boundary conditions are implemented at *r* = 0 μm and *r* = 4350 μm. The numerical solutions of Eq. (1)-(2) are obtained with Δ*r* = 10 μm Δ*t* = 0.1 hours and *ε* = 1 × 10^−5^.

Results in Fig. 4-5 indicate that cell density profile for co-culture 1 appears to contain a ‘dip’ in the central region, near *r* = 0 μm. Equivalent results for co-cultures 2, 3, and the monoculture experiments do not contain such a pronounced dip (Fig. 4-5). Since all experiments are prepared using the same procedure, we prefer not to offer an interpretation of this dip because it could be due to a statistical fluctuation rather than some underlying mechanism.

All of our results, so far, suggest that the diffusivity and proliferation rate of primary fibroblast cells and SK-MEL-28 melanoma cells are insensitive to whether the cell populations are grown in monoculture or co-culture. This implies that there is limited interactions or crosstalk between these two cell populations. If we were to observe some interactions, such as melanoma cell migration being stimulated by the presence of fibroblast cells, we can use our mathematical model to explore how these potential interactions might be best observed. To explore this we conduct a series of numerical experiments to investigate how the solutions of Eq. (1)-(2) depend on *D_SK_*. Since we focus on altering *D_SK_* alone, we find that the solution of Eq. (1)-(2) is most sensitive at the low-density leading edge of the population, as depicted in Fig. 6(a). Results in Fig. 6(b) compare the experimental density profile for co-culture 3 at *t* = 48 hours with the standard choice of parameters from Fig 5, showing that the solution of the mathematical model for *S*(*r*,*t*) matches the experimental data quite well at the leading edge of the spreading population. We also show results where *D_SK_* is increased by a factor of 5, where we see that the solution of the mathematical model predicts that the population spreads notably further than observed in the experiments. Similarly, we also show equivalent results where *D_SK_* is increased by a factor of 20 and the differences are even more pronounced.

**Fig. 6.**
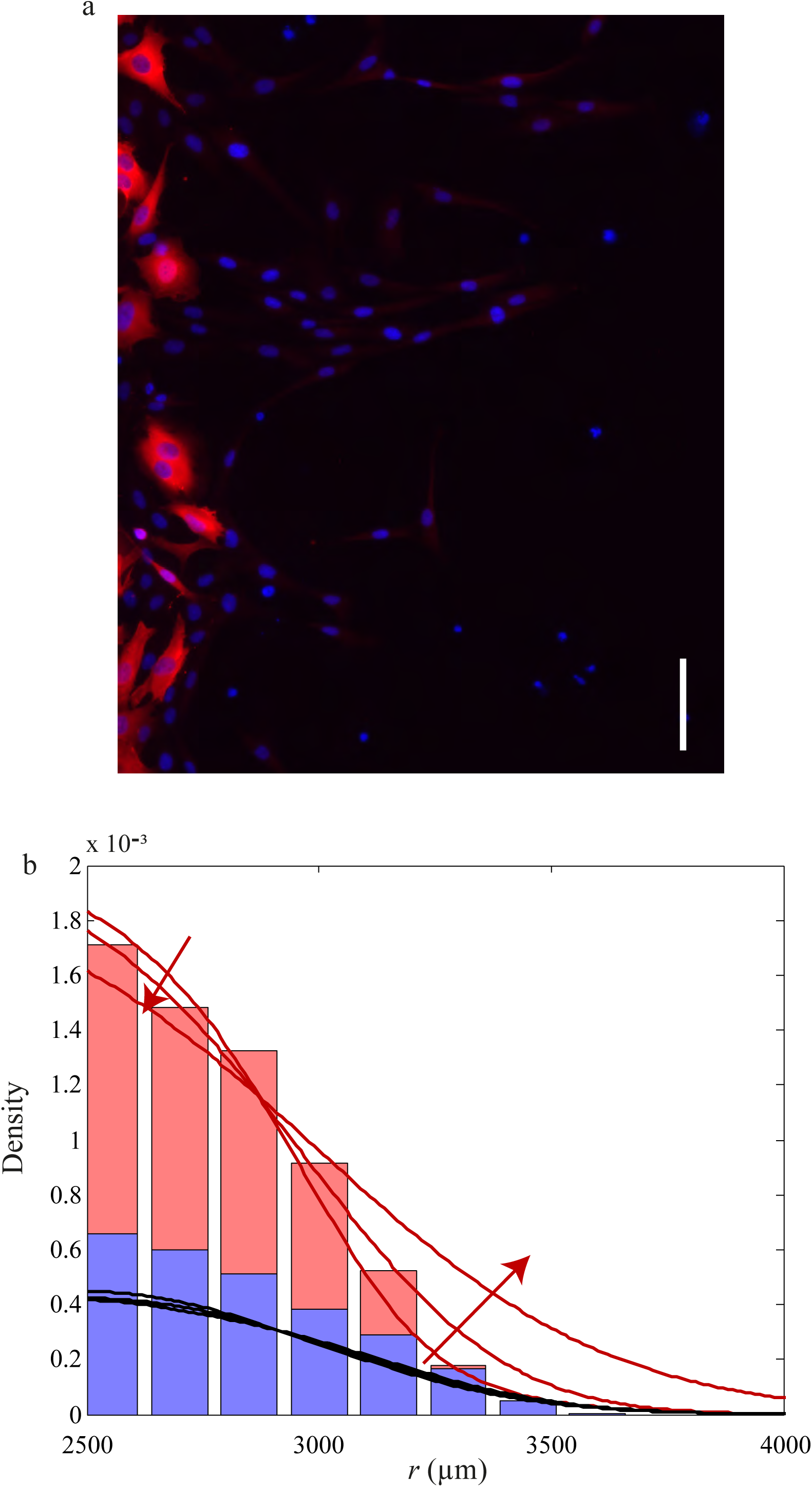
Focussing on the leading edge of the spreading population highlights the sensitivity of the spreading to the value of *D_SK_*. (a) Immunofluorescence staining at the leading edge of the spreading population in co-culture 3 showing individual fibroblast cells (blue) ahead of the SK-MEL-28 melanoma cells (red). The scale bar is 100 μm. (b) The average experimental cell density profile for co-culture 3 after 48 hours near the leading edge of the spreading population is compared with three numerical solutions of Eq. (1)-(2). The solution for *C_Fb_* (*r*,*t*) is shown in black, *S*(*r, t*)is shown in red and *C_SK_* (*r, t*)is the difference between the red and black curves. The parameters used to solve Eq. (1)-(2) are: *D_Fb_* = 1200 μm^2^/hour; *λ_Fb_* = 0.03 /hour; *λ_SK_* = 0.03 /hour and *K* = 2.8 × 10^−3^ cells/μm^2^ for all three profiles, while the value of *D_SK_* includes 170, 850 and 3400 μm^2^/hour with the arrows showing the direction of increasing *D_SK_*. The equations are solved on 0 < *r* < 4350 μm, but the profiles are shown only at the leading edge where 2500 < *r* < 4000 μm. The numerical solutions of Eq. (1)-(2) are obtained with Δ*r* = 10 □m, Δ*t* = 0.1 hours and *ε* = 1 × 10^−5^.

The additional results in Fig. 6 suggest that the comparison of the experimental cell density profile and the solution of the mathematical model at the leading edge of the spreading population is relatively sensitive to the value of *D_SK_*. One way of interpreting the results in Fig. 6(b) is that our procedure could be used to detect potential interactions that would lead to a modest increase in *D_SK_* when the two cell types are grown in co-culture.

## 6. Conclusion

Many *in vitro* studies examining the spatial spreading of cancer cells use monoculture experiments (Kramer et al., 2013; Liao et al., 2013; Treloar et al., 2013; Treloar et al., 2014b). However, these monoculture experiments are unrealistic in the sense that the spreading cancer cells are not subject to interactions with other cell types as would occur *in vivo*. To address this limitation, our approach to investigate the spatial spreading of a melanoma cell population is to examine a suite of monoculture and co-culture circular barrier assays. Our monoculture experiments involve studying both primary fibroblast cells and SK-MEL-28 melanoma cells separately, while our co-culture experiments use three ratios of both cell types in the same experiment. All of our experiments are initialised by placing approximately 20,000 cells into the barrier, and we examine the collective spreading over a period of 48 hours. To quantify how these heterogeneous populations of cells spread over time we: (i) measure the diameter of each expanding cell population; (ii) identify individual cell types in each expanding population; (iii) count the number of primary fibroblast cells and SK-MEL-28 melanoma cells in a series of transects across each expanding cell population; and (iv) construct cell density histograms to show the spatial arrangements of each cell type in each expanding cell population.

Previous experimental studies suggest that interactions between fibroblast and melanoma cells can lead to an increase in rates of collective spreading when these two cell types are in contact (Cornil et al., 1991; Flach et al., 2011). However, these previous studies do not make detailed measurements of spatial and temporal arrangements of cells within the spreading population, and they do not consider varying the initial ratio of fibroblast cells to melanoma cells. To provide more information about the interaction between melanoma cells and fibroblast cells, we perform co-culture experiments using three different initial ratios of cells, and we make detailed measurements about the spatial and temporal arrangements of both subpopulations within the heterogeneous population as it spreads and grows. In summary, our experimental results indicate that the influence of primary fibroblast cells on SK-MEL-28 melanoma cell growth and spatial expansion in a two-dimensional circular barrier assay is minimal. To provide additional information about this apparent lack of interaction, we also calibrate a mathematical model to our experimental data.

In this work, we use our mathematical model to provide a novel analysis of experiments exploring potential interactions between fibroblast cells and melanoma cells. By first estimating the cell diffusivity and cell proliferation rate for the primary fibroblast cells and SK-MEL-28 melanoma cells separately in monoculture, we find that the parameter estimates are consistent with previously published estimates for mouse fibroblast cells and other metastatic melanoma cell lines (Treloar et al. 2013; Treloar et al. 2014a). Therefore, we are confident in our estimates of the cell diffusivity and cell proliferation rate in monocultures because they are consistent with previously published data for similar experiments. Then, to address the question of whether cells in monoculture behave similarly, or differently, to cells in co-culture, we investigate whether the solution of the co-culture mathematical model, parameterised using estimates from the monoculture experiments, can genuinely predict the behaviour of the co-culture experiments. Since we find that a fixed choice of parameters, that is very similar to the estimates from the monoculture experiments, can predict both the spatial and temporal patterns of collective spreading in the two monoculture experiments, and all three co-culture experiments, we conclude that the spreading and growth patterns observed for primary fibroblast cells and SK-MEL-28 melanoma cells are not affected by growing them in monoculture or co-culture.

Our approach is to compare the spatial spreading of two different cell types in both monoculture and co-culture circular barrier assays to quantify the rates of cell migration and the rates of cell proliferation for each cell type. This approach is novel because most combined experimental and mathematical modelling studies focus on monoculture experiments alone (Kramer et al., 2013; Treloar et al., 2013). However, it is possible to explore other alternative experiments to further extend our work. This includes performing additional experiments to examine the rates of spatial spreading in different cell lines. For example, it could be of interest to repeat our work using other kinds of melanoma cell lines including cell lines from earlier stages of the disease, such as melanoma cells associated with the radial growth phase or the vertical growth phase (Haridas et al., 2016). This could be an important consideration because our current work focuses on examining potential interactions between fibroblast cells and SK-MEL-28 melanoma cells only. Since the SK-MEL-28 cell line is associated with the metastatic phase (Fofaria and Srivastava, 2014; Haridas et al. 2016), it is possible that these cells have progressed beyond being influenced by fibroblasts. Alternatively, if our experiments and analysis were repeated using melanoma cells from earlier stages, it is conceivable that these melanoma cells might be more responsive to the presence of fibroblasts.

Other options to extend our work might involve incorporating further types of cells to make the co-culture experiments more realistic. For example, it is of interest to include both primary fibroblast cells and primary keratinocyte cells in a co-culture experiment with melanoma cell lines. However, this kind of extension is difficult because we would need to specifically identify three different cell types to understand how the three different subpopulations are spatially arranged. Therefore, we leave this extension for future consideration.

## Acknowledgments

We acknowledge financial support from the Australian Research Council (FT130100148, DP140100249), and we thank Professor Brian Gabrielli for providing us with the SK-MEL-28 melanoma cell line. Helpful suggestions from the two referees are appreciated.

